# A VTA GABAergic computational model of dissociated reward prediction error computation in classical conditioning

**DOI:** 10.1101/2020.02.06.936997

**Authors:** Pramod Kaushik, Jérémie Naudé, Surampudi Bapi Raju, Frédéric Alexandre

## Abstract

Classical Conditioning is a fundamental learning mechanism where the Ventral Striatum is generally thought to be the source of inhibition to Ventral Tegmental Area (VTA) Dopamine neurons when a reward is expected. However, recent evidences point to a new candidate in VTA GABA encoding expectation for computing the reward prediction error in the VTA. In this system-level computational model, the VTA GABA signal is hypothesised to be a combination of magnitude and timing computed in the Peduncolopontine and Ventral Striatum respectively. This dissociation enables the model to explain recent results wherein Ventral Striatum lesions affected the temporal expectation of the reward but the magnitude of the reward was intact. This model also exhibits other features in classical conditioning namely, progressively decreasing firing for early rewards closer to the actual reward, twin peaks of VTA dopamine during training and cancellation of US dopamine after training.

## Introduction

The phasic firing activity of midbrain dopamine neurons is believed to encode a reward prediction error, which can guide learning and serve as an incentive signal. In his famous experiment, Pavlov observed that if food follows the ring of a bell, a dog comes to salivate after the bell is rung. This process is called classical (or pavlovian) conditioning: an unconditioned response (salivation) originally associated with an *Unconditioned Stimulus* (US, the food) becomes conditionally elicited by a *Conditioned Stimulus* (CS, the bell ring). Schultz and collaborators examined the activity of midbrain dopamine neurons in primates, during a classical conditioning task similar to Pavlov’s. They observed that originally, midbrain dopamine neurons responded with a burst of spikes to unexpected primary rewards (juice/water dripped in the mouth of thirsty primates), i.e. an US. After the primates learned the association between a tone CS and the reward US, dopamine cells responded to the reward-predicting CS. Moreover, these neurons stopped responding to US whose arrival was expected, being predicted by the CS. When cued rewards failed to be delivered, a large percentage of dopamine neurons revealed a brief pause with respect to their background firing at the moment of expected reward in Figure 2.

**Figure 1.**
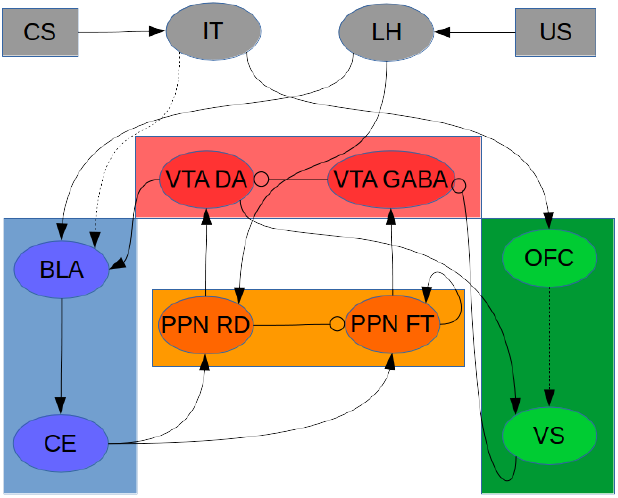
Model diagram illustrating the neuronal structures and their connections involved in Reward Prediction Error (RPE) computation. Pointed arrows represent excitatory connections, while rounded arrows represent inhibitory projections. Dashed lines represent learnable connections, while solid lines represent fixed connections in the model. Ventral Tegmental Area has Dopamine (DA) and GABA populations which play a central role in computing RPE. This RPE is transmitted to Basolateral Amydala (BLA) and the Ventral Striatum (VS) where the modulatory connections undergo change depending on this signal. Both the VTA structures receive inputs from distinct Peduncolopontine (PPN) neural populations. The PPN Reward Delivery (RD) neurons deliver reward to VTA DA neurons from the Lateral Hypothalamus (LH) which receives the unconditioned stimulus (US). The PPN Fixation target (FT) neurons receive their projections from the Central Nucleus (CE) of the Amygdala and these PPN neurons along with Ventral Striatum (VS) project to VTA GABA forming the inhibitory signal that cancels the VTA DA upon reward delivery. The BLA is regarded to learn the association between unconditioned stimulus (US) and the conditioned stimulus (CS) and produces the anticipatory firing in VTA DA through the BLA->CE->PPN RD->VTA DA pathway. The VS is posited to learn the timing of the interval and it has inhibitory projections on VTA GABA.

**Figure 2.**
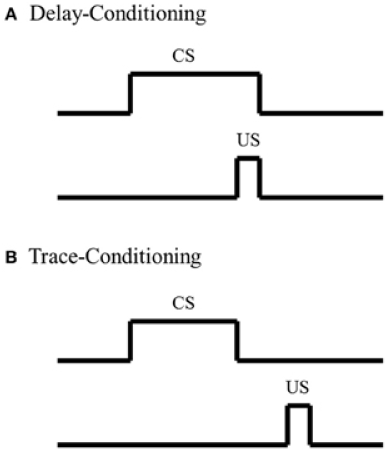
Delay Conditioning - In this paradigm, the conditioned stimulus (CS) persists until the presentation of the unconditioned stimulus (US)

This predictive behavior of VTA Dopamine neurons was linked to the *Temporal Difference Learning* algorithm in Reinforcement Learning (***Sutton and Barto, 1998***). The TD algorithm predicts a future reward that occurs at a specific state ahead of time after a given number of trials. This prediction is arrived through the computation of a reward prediction error (RPE) that enables learning the value of each state. It is represented by the equation:

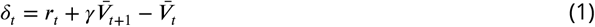

where *r_t_* is the reward (return) on time step *t*. Let 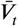 be the correct prediction that is equal to the discounted sum of all future rewards. The discounting is done by powers of factor of *γ* such that reward at distant time step is less important as compared to recent rewards.

The TD algorithm is powerful in its heuristic value, as it links the activation of dopamine cells with classical conditioning. However, some properties of TD models are inconsistent with the electrophysiological data, and it lacks the biological realism needed to explain the mechanisms through which RPE-like activity emerges in dopamine cells. Most inconsistencies are concerned with the way time is represented in the TD model. Whereas the TD error signal travels back in time from US to CS when learning progresses, it has been reported in (***Schultz et al., 1997***) that US-related activation of VTA slowly decreases while the CS-related one increases. Ramping dopamine activity during the CS-US interval has been proposed to reflect back-propagating error signals averaged over trials (***Niv et al., 2005***) but this ramping can be observed in individual trials and is actually related to reward uncertainty (***Fiorillo et al., 2003***)(***Fiorillo et al., 2005***). More importantly, when US is delivered earlier than predicted, VTA dopaminergic neurons are activated at the actual time of the US but not at the usual time of reward (***Hollerman and Schultz, 1998***), contrarily to what is predicted by TD. Critically, when striatum is lesioned, experiments (***Takahashi et al., 2016***) show that VTA dopaminergic neurons signal a RPE when reward magnitude changes, but not when time of the reward is modified. This suggests that learning about reward timing is computationally and anatomically separated from learning about reward magnitude, i.e. a completely different implementation of reinforcement learning than is usually considered ***Joel et al., 2002***).

Here we sought a system-level account of how the CS-US interval duration and the value of the reward are separately learned, and how these two features are combined to give rise to dopaminergic phasic activity. In particular, we considered two components of the meso-limbic loops that have been overlooked in previous models: VTA GABA cells and neurons from the Pedunculopontine Nucleus (PPN). We propose that VTA GABA neurons provide inhibitory drive onto VTA DA neurons (***Eshel et al., 2015***), display persistent firing during the CS-US interval (***Cohen et al., 2012***) and are necessary to compute the RPE in DA cells (***Eshel et al., 2015***). Neurons from the PPN project to both VTA DA and GABA neurons and have been found necessary for appetitive conditioning (***Yau et al., 2016***). PPN neurons are classically believed to signal the delivery of actual reward (***Vitay and Hamker, 2014***). However, during conditioning, two types of response are recorded from neurons from the PPN: neurons responding to the actual reward magnitude, and a persistent neuronal firing during the CS-US interval, reflecting the prediction or expected reward magnitude (***Okada and Kobayashi, 2009***)(***Okada and Kobayashi, 2013***). Here we show that a system-level computational model can account for these yet-unexplained physiological and lesion data. We propose that the Ventral Striatum learns the reward timing and the Amygdala the reward magnitude, which is then transferred to the PPN. Expected reward timing and magnitude are subsequently combined at the level of VTA GABA cells to compute the expectation term needed by VTA DA neurons to generate a prediction error. We furthermore provide testable predictions for future experiments, on the role of PPN and VTA GABA in classical conditioning.

## Results

We built a system-level network in which reward prediction error (RPE) emerges during learning, from the interaction of the neruronal ensembles repersenting RPE computation. To illustrate how RPE is computed in the network, we subjected it to in silico experimental scenarios similar to a conditioning task. This analogue of classical conditioning consists in the repeated pairing of a CS with the US, separated by a fixed interval duration. The in silico experiments below examine how the value (magnitude) of the US reward and the duration of the CS-US interval are learned separately, and combined by VTA GABA cells. In turn, VTA GABA neurons provide the expectation term used to cancel the excitation of dopamine cells by the US, at the time of the expected reward (***Cohen et al., 2012***)(***Eshel et al., 2015***). Finally, we show how this new model provides a better description of experimental data that have been left unexplained yet, i.e. how dopamine cells respond to rewards delivered earlier than expected (***Fiorillo et al., 2008***), and how value and timing are dissociated by VS lesions (***Takahashi et al., 2016***).

### Model Architecture

The system-level computational model attempts to explain how the dopamine reward prediction error is computed in appetitive conditioning in the VTA through understanding the roles of VTA GABA and Peduncolopontine (PPN) neurons. The model is shown in Figure 1. The model focuses on the computation inside the VTA carried out by two populations viz., the VTA DA and VTA GABA neurons. Lateral Hypothalamus (LH) projects to PPN RD and VS and these in turn project to VTA DA and VTA GABA respectively. When reward is delivered, it is reported to fire the Lateral Hypothalamus (LH) and activates the LH → PPN RD (Reward delivery) → VTA Dopamine pathway resulting in US dopamine firing prior to any sort of learning (***Semba and Fibiger, 1992**; **Lokwan et al., 1999***). Basolateral Amygdala (BLA) learns the magnitude of the US through the projections from VTA DA to BLA, which signal a reward prediction error that modulates synaptic plasticity. A pathway from LH to BLA learns that BLA firing for CS has the same amplitude as the US-induced firing (***Sah et al., 2003***). The immediate firing in response to the CS occurs through the BLA recognizing the cue encoded by the Infero-temporal cortex (IT), with BLA activating the VTA DA through the BLA→ CE → PPN RD → VTA DA pathway. The Central Nucleus of the Amygala (CE) has excitatory projections on PPN FT(Fixation Target) (***Okada and Kobayashi, 2013***) (***Kobayashi and Okada, 2007***) which displays persistent activity unless strongly inhibited. The LH also projects to the VS, which is also modulated by VTA DA neurons. Finally, excitatory projections from PPN FT and inhibitory projections from VS neurons to VTA GABA enable the final reward cancellation of VTA DA observed in electro-physiological experiments.

### Control Scenario

In the following, we show how a reward prediction error emerges in VTA Dopamine cells with learning, as a consequence of network dynamics and plasticity.

- Initial Trial The arrival of an unexpected reward induces a firing in the LH neurons. This LH firing subsequently activates VTA Dopamine neurons, through the PPN RD neurons (Figure 3 B top panel Trial1, Figure 3 C top panel Trial 1). Hence, the model reproduces that VTA Dopamine neurons fire upon the delivery of an unexpected reward. No activity in BLA and Ventral Striatum (VS) and VTA GABA (Figure 3 B bottom panel Trial1) is observed at this stage of learning. Indeed, Basolateral Amygdala (BLA) has not yet learned to associate the magnitude of the CS with the US, and the Ventral Striatum (VS) the timing of the CS-US interval duration. Hence, there is no expectation at the arrival of the CS.
- Partial Conditioning The synaptic weights between IT and BLA are updated after each rewarding trial. Consequently, after a few trials (7 in our simulations), the BLA starts responding to the CS stimulus. This progressive learning in the BLA generates firing in VTA DA (Figure 3 B top panel) through PPN RD (Figure 3 C top panel) in response to the arrival of the CS, corresponding to a partial prediction of reward. BLA activity also generates a partial expectation through tonic firing in PPN FT (Figure 3 C bottom panel Trial 7). Hence, a partial cancellation of VTA dopamine neurons happens at this stage. At this stage, the time interval has been completely learned, contrary to the learning of the US magnitude. This corresponds to the activity of the Ventral Striatum reaching its minimum at the exact moment of reward. Hence, the VS does not exert any inhibition at the end of the interval, which results in the inhibition of VTA DA neurons at the expected time of the US. However, as the US magnitude is not fully learned yet, the activity of PPN FT to VTA GABA pathway only results in partial cancellation of VTA DA activity upon US delivery. This is consistent with experimental results on Partial conditioning (***Pan et al., 2005***), and designed as a partial expectation in the TD framework. (Figure 3 B bottom panel Trial 7). This partial expectation consists in a twin peak of VTA firing, at the respective times of the CS and the US.
- Complete Reward Cancellation The final state of the circuit, after 16 CS-US pairings, is given in Figure 3 A. The magnitude of expectation originates from the CS firing in the Central Amygdala (CE) and is maintained in the PPN FT through a self-sustaining mechanism (***Okada and Kobayashi, 2013***) (***Kobayashi and Okada, 2007***). At the end of learning, the BLA neurons have reached an asymptote in their firing, which encodes for reward magnitude. Hence, PPN FT neurons display maximum tonic activity after the presentation of the CS. In parallel, as in partial conditioning, the presentation of the CS, which fires the IT and thereby the Orbitofrontal Cortex (OFC) (***Carmichael and Price, 1995***), activates the VS that encodes the interval timing. It acts similar to a negative integrator and progressively lowers the inhibition that VS exerts on PPN FT, to reach zero inhibition at the expected time of the reward. Both signals are combined by VTA GABA, the activity of which peaks at the time of the reward (Figure 3 B bottom panel Trial 16), cancelling VTA dopamine, which no longer shows firing at the time of the US when reward arrives through the LH (Figure 3 B top panel Trial 16). The magnitude of expectation originates from the CS firing in the Central Amygdala (CE) and maintained in the PPN FT through a self sustaining mechanism (***Okada and Kobayashi, 2013***) (***Kobayashi and Okada, 2007***). The GABA firing in the VTA is reflective of this (***Yau et al., 2016***) and the PPN FT integrates the magnitude from the Central Amygdala (CE) and timing information from the VS to achieve the ramping signal that encodes both time and magnitude of the reward delivery.

**Figure 3.**
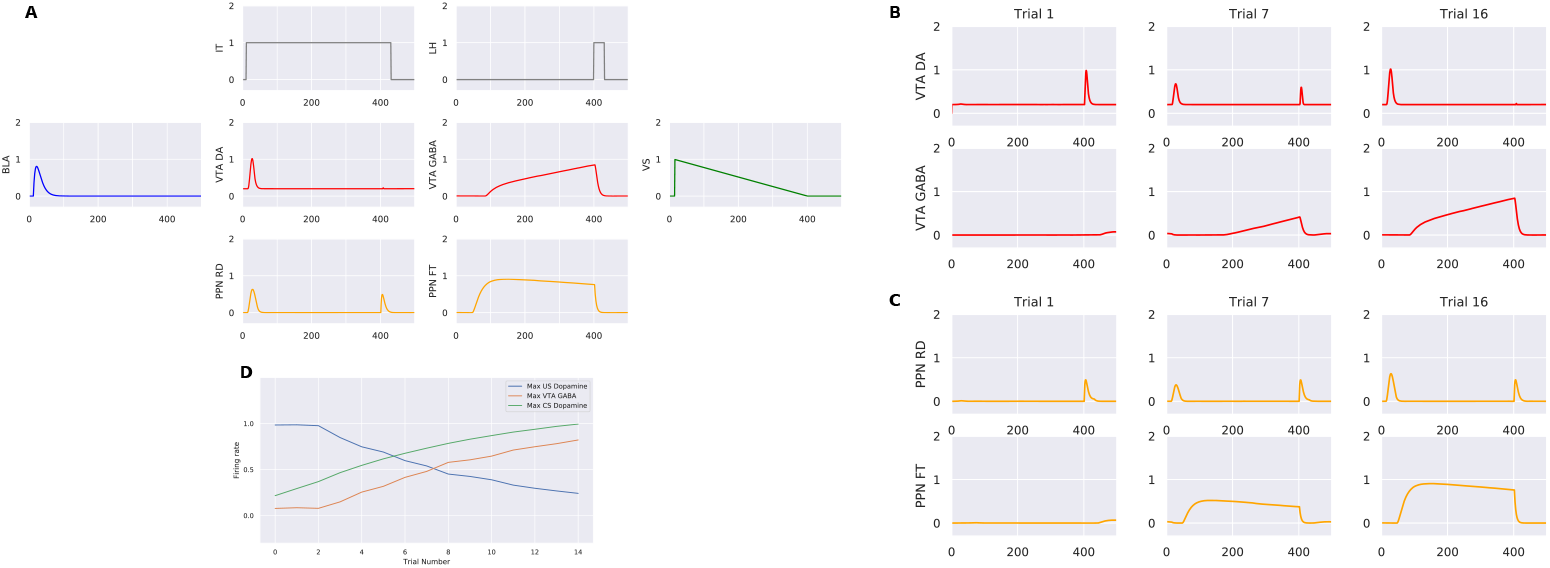
(A) The figure represents the firing of the neuronal populations after training following the same color conventions as the model architecture diagram. IT and LH indicate the stimulus and the reward respectively. PPN RD represent the phasic signals received from BLA and the LH at stimulus onset and reward delivery. VTA DA neurons show firing only at the arrival of the CS and VTA GABA neurons have a ramping expectation signal that fully cancels the VTA DA signal at reward delivery. BLA and VS have encoded magnitude and timing of the reward respectively. (B) This figure portrays the evolution of the VTA and PPN sub-populations throughout the duration of trials. VTA DA shows firing only at the reward delivery during initial trials while VTA GABA shows no firing due to absence of any expectation at this stage. At Trial 7, VTA DA shows twin peaks reflecting both a partial encoding at stimulus arrival and partial cancellation of reward signal. VTA GABA on the other hand has a ramping nature at the precise time of the arrival of the reward indicating encoding of timing and partial encoding of expectation. The final trial diagram shows the complete cancellation of the VTA DA signal at reward delivery and a complete encoding of stimulus at the CS arrival. Correspondingly, VTA GABA has a larger expectation signal that ramps that has completely encoded the magnitude of the reward signal. (C) shows the evolution of the PPN sub-populations during the sequence of trials with PPN RD signalling reward delivery from LH to VTA DA during the initial trials and PPN FT not showing any sign of expectation at the same time. In the middle of training, PPN RD reflects a partial firing for stimulus onset and the same reward signal firing as during the initial trials. PPN FT also reflects this partial firing with a tonic nature of firing that passes onto VTA GABA. During the final trial stages, PPN RD firing peaks for its stimulus arrival firing while a bigger tonic firing signal is observed for PPN FT. (D) portrays the evolution of VTA DA CS and US signals and that of VTA GABA. Both VTA GABA and VTA DA CS show increased firing across the trials while VTA DA US progressively reduces ultimately to the background firing rate of VTA DA. All the figures are averaged for 10 runs.

### Variability in Magnitude and Time

1. Variability in Timing When a reward is delivered earlier than expected (i.e. with a shorter delay than the CS-US interval that has been learned), US firing is observed in VTA dopamine neurons. More precisely, in this case of earlier-than-expected US reward, VTA dopamine neurons fire less than the initial (before learning) firing observed at US delivery. This is consistent with the US being expected, albeit not at this precise timing. An interpretation would be that partial expectations are generated during the CS-US interval, hence the reward prediction decreases with time until the expected timing of US delivery. In our model, we observed that the earlier the reward was delivered, the higher was the VTA DA firing (Figure 4 B). VTA DA firing in Figure 4 A (middle left panel) shows a reward delivered before the half-way-point (at the 100^*th*^ time step) evoking a dopamine firing, but less than the firing initially induced by an unpredicted reward. By contrast, Figure 4 B (middle left panel) shows an early reward delivered after the half-way-mark (at the 300^*th*^ time step), which induces lesser firing in VTA DA cells. The model is consistent with physiological data in primates, where earlier-than-expected rewards evokes progressively less firing as the reward delivery time increases (***Fiorillo et al., 2008***). Our model provides a mechanistic explanation for this data.
2. Variability in Magnitude VTA Dopamine firing reflects the difference between the actual reward and the expected reward. For instance, VTA Dopamine neurons fire on US arrival if a reward is larger than expected (Figure 5 C middle panel). This corresponds to a positive reward prediction error, in accordance with physiological data. The subtractive nature of inhibition (***Eshel et al., 2015***) encodes the difference of magnitude between the actual magnitude of reward and the expected magnitude of reward (Figure 5C middle panel).

**Figure 4.**
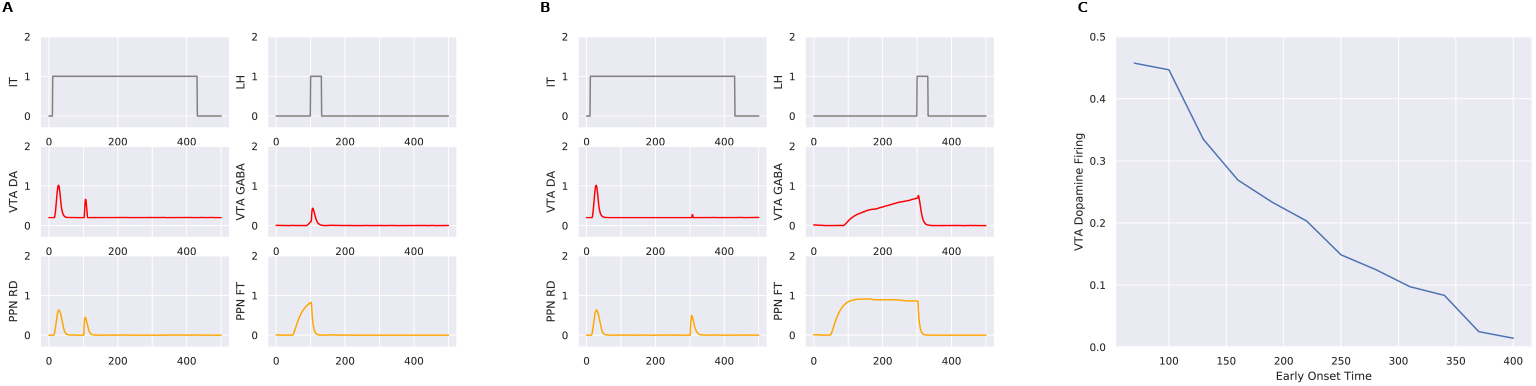
These figures indicate the early firing scenarios on VTA and PPN sub-populations. LH indicates the reward delivered earlier than usual and IT the stimulus. (A) denotes the arrival of the reward at 100^*th*^ time step after training where VTA DA shows some firing compared to no firing after training when reward is delivered at the usual time. (B) This firing for an earlier reward is still larger than a later arrival of early reward at 300 time steps where VTA DA barely shows any firing. (C) indicates the progressively later “early” rewards fire less as early reward delivery times get closer to usual reward arrival times in accordance with data on (***Fiorillo et al., 2008***)

**Figure 5.**
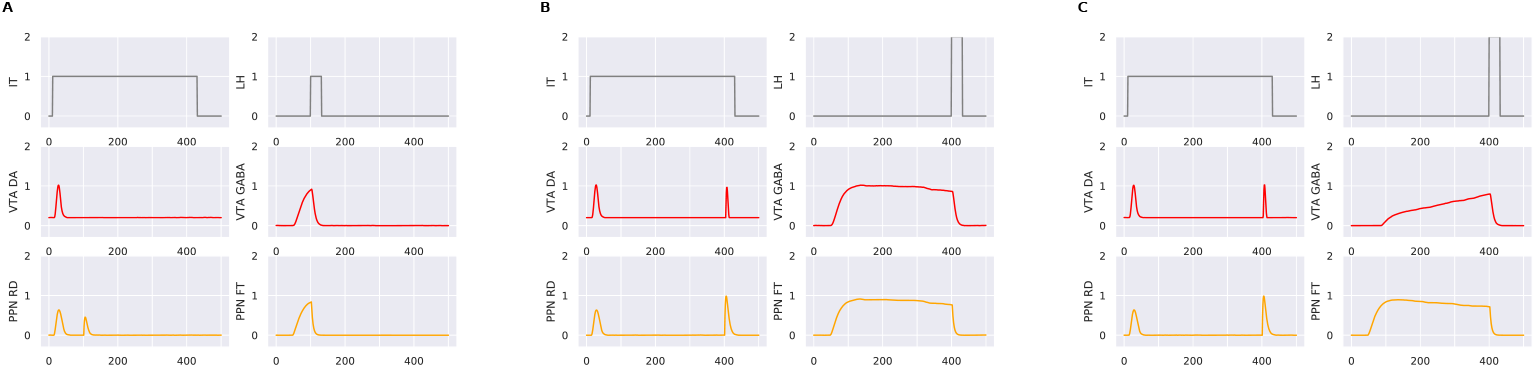
Panel3-These figures indicate how timing information is lost but magnitude information of the reward is maintained on VTA sub-populations. LH indicates the reward delivered earlier than usual and IT the stimulus. During the VS lesion scenario, VTA GABA is no longer able to integrate the timing information coming from VS and just reflects the projection from PPN FT neurons. Hence, early rewards are treated as any other reward that comes at the usual time, reflecting no firing (left panel). But when the reward is of a higher magnitude (middle panel), it is able to indicate the prediction error just as in the control scenario where VS is not lesioned and VTA GABA integrates both magnitude and timing.

### Dissociation of time and magnitude prediction errors following VS Lesions

1. VS lesion affects time prediction error When an experimental lesion is made in the VS from rats (***Takahashi et al., 2016***), earlier-than-expected US does not trigger firing in VTA Dopamine cells. Accordingly, our model reproduces this lack of timing prediction error, as the virtual lesion of the VS in the model abolishes the VTA Dopamine response to earlier-than-expected US. Moreover, VTA GABA cells, instead of displaying a ramping signal, provide a constant tonic inhibition, similar to PPN FT, throughout the duration of the trial (Figure 5 A middle right panel). Indeed, the model posits that lack of dopamine firing for the VS-lesioned scenario compared to the control scenario is due to the higher inhibition from the VTA GABA neurons (Figure 5 A middle right panel). The VS inhibitory signal acts as a “when” signal providing VTA GABA neurons with information enabling computation of reward expectation at a given point in time. Due to this information being lost because of lesion, VTA GABA neurons do not have a ramping signal that peaks at the right moment. Instead, they have a constant inhibition throughout the interval.
2. VS lesion does not alter magnitude prediction error The rewards with higher magnitude induce firing in VTA DA cells even if the VS is lesioned. A reward that is double the magnitude triggers the same effect on US VTA Dopamine as in the control scenario (Figure 5 B middle left panel). Since the VS encodes only temporal information, its lesion does not affect the magnitude encoding of the stimuli. This results in VTA DA firing showing the same subtractive effect as in the control scenario(Figure 5 B middle left panel). This is due to the VTA GABA encoding the magnitude from the BLA through the PPN FT neurons. A lack of VTA GABA ramping does not prevent the VTA DA from having the magnitude encoded within its population, and provides the same RPE at usual reward time. This behavior of VTA DA neurons in the case of a VS lesion with larger magnitude is also in accordance with experimental results (***Takahashi et al., 2016***).

## Discussion

### Role of Timing in conditioning

The importance of CS-US interval timing is one of the key postulates of the proposed theory underlying the model. In this model, the interval timing mechanism uses the dopamine signal since the onset of the CS to learn the underlying temporal distribution to predict the arrival of the reward, however it needn’t be the case. Learning the interval time separating the CS and the reward could happen without dopamine with a phenomenon called sensory-preconditioning (***Sadacca et al., 2016***). Our model predicts that since time and magnitude are separate signals in the brain, the learning of time precedes the learning of magnitude for reward prediction error to take place. Interval timing learning in animals has been observed to happen in very few trials and sometimes even one. The model posits that interval timing learning is an integral part of reward prediction error computation in appetitive learning and the learning of timing happens before the magnitude of the stimulus so as to construct an inhibitory signal that has a ramping nature before it ramps to its maximum amplitude. The timing mechanism used in the model is very simplistic, reflecting the learning of a single parameter to get the final slope of inhibition. It is possible that more complex timing mechanisms are incorporated in the striatum to accommodate stochasticity in the temporal interval.

This is consistent with the original Temporal Difference Reinforcement Learning (TDRL) representation of dopamine where the cue is tracked since the onset of its presentation, state by state until reward is delivered as in the complete serial compound (CSC) stimulus representation (***Schultz et al., 1997***). Here too, the state is tracked at each time point using the timing signal. However, unlike the TD model, which is model-free and tracks value across states, this model learns a separate value signal for state similar to the Successor Representation (SR)(***Dayan, 1993***) where reward and state representation are computed separately unlike TD-Learning. This dissociation enables the model to learn changes in magnitude independent of the state and learn the state independent of the magnitude.

### Dual pathway model

The dual pathway models of classical conditioning posit that the mechanism with which the CS firing occurs at the onset of a trained cue is dissociated from the expectation that inhibits the reward signal at the time of the reward. Our model too, largely follows the same pattern but with some deviations (***O’reilly et al., 2007***). This model replicates the observation in ***Fiorillo et al*.** (***2008***) about early rewards delivered at different time points exhibiting different firing. The interpretation of this model is that, this dopamine error is indicating a mismatch in state rather than a mismatch in magnitude. Since magnitude is a separate signal and does not require updating, the early reward firing is indicative of a violation of a belief of the state the animal is in. This prediction error is predicted to happen between the striatal inhibition and VTA GABA. This inhibition is also predicted to play a role in the lesion experiments done by ***Takahashi et al*.** (***2016***) where VS lesions hamper the ability of the animal in tracking state. This lesion inturn has an effect on VTA GABA which is no longer inhibited and only serves the magnitude component of the reward prediction error. Hence, early reward prediction errors no longer happen, because the animal does not recognize state prediction errors and expects the reward to happen all the time. It was also found in the study that VTA non dopamine neurons have a higher firing after the VS lesions consistent with our model and our model predicts, VTA GABA to have higher firing when VS is lesioned. This model hypothesis a continuous ramping signal that is active throughout the duration of the CS and the US, peaking at the time of the US and speculates that this inhibition signal and the CS firing could have the same source.

### Heterogeneity of PPN

Though much of PPN’s anatomical and chemical characteristics are unknown, studies have shown functional differences between populations inside PPN. Specific populations within PPN exist which fire phasically for rewards and others which have sustained firing from reward prediction to delivery of reward, and the activation sustaining till the delivery of the reward even if the arrival of the reward is delayed (***Okada and Kobayashi, 2009***). Moreover these neurons seem to encode amount of reward firing higher for rewards with larger magnitude (***Okada and Kobayashi, 2009***) (***Hong and Hikosaka, 2014***) portraying a graded firing signal capable of differentiating reward amounts consistent with this models characteristics. Studies have also pointed out a growing role for PPN projections to VTA non-dopamine neurons as necessary for appetitive pavlovian conditioning (***Yau et al., 2016***) and activation of PPN glutamate neurons to be reinforcing (***Yoo et al., 2017***). The hypothesis of this model is that a subset of PPN neurons (PPN FT) convey reward magnitude information to the VTA GABA neurons and any optogenetic silencing of these neurons can interfere with the computation of reward prediction error, though isolating these neurons could prove difficult owing the structures heterogeneity.

### VTA GABA theory of computing RPE in classical conditioning

Previous studies have implicated the striatum (***Usuda et al., 1998***) as the source of the inhibitory signal cancelling the dopamine. But, recent projection-specific activation by optogenetic studies among others have shown that the inhibition from striatum has weak to no inhibitory effects on DA neurons when stimulating direct striatal inputs on DA neurons in the VTA (***Keiflin and Janak, 2015***) (***Bocklisch et al., 2013***) (***Chuhma et al., 2011***)(***Xia et al., 2011***)(***Klein-Flügge et al., 2011***). There have been a number of recent results, suggesting an alternate pathway within VTA which might be responsible for this inhibitory signal. Moreover, optogenetic studies done on VTA (***Cohen et al., 2012***)(***Eshel et al., 2016***) have pointed out not only does the VTA GABA neurons exert enough inhibition to cancel VTA DA neurons but the inhibition is also subtractive in nature and hence suitable for computation of reward prediction error (***Eshel et al., 2015***). This model hypothesises that the ramping nature of expectation which encodes both magnitude of the stimulus and time of arrival is encoded in the VTA GABA which acts as the site of integration between these two different dimensions of the reward. One possible explanation of this distributed nature of reward prediction error could be that this is what allows for rapid recomputation of values (either of time or magnitude) and allows the animal to exhibit and sometimes fast, adaptive behavior. Parallels could be drawn with the literature of reinforcement learning that the animals are not purely engaged in model-free reinforcement learning and that the dopamine signal itself could be not just performing reward prediction errors and differences in timing could elicit an error from the dopamine system for state prediction errors. For example, dopamine firing for early reward delivery could be interpreted as a state prediction error where the animals has to reevaluate the time of the reward rather than the magnitude of the reward. The precise interpretation of the dopamine prediction error could be handled by the upstream areas to determine what computations are to be done to reflect a changed scenario.

This model examines the role of VTA GABA in computing the reward prediction error along with a few other subtrates based on some of the results provided by ***Takahashi et al*.** (***2016***). This paper adoptsy a semi-markov approach to explain the findings while the model given here attempts to provide a system level model of how the underlying neuronal substrates might act. There are a few other behaviors that is observed in the model. The authors note that removing VS does not remove expectation and the animal in effect expects reward all the time, this could be the VTA GABA signal we observe in the model when VS is lesioned. VTA GABA loses its ramping functionality and has a flat tonic firing pattern carrying on its earlier peak expectation thoughout the duration of the trial, The authors also note that “non-dopaminergic” neurons show significantly higher baseline firing rate when VS is lesion. Our model hypothesises that it is indeed the VTA GABA neurons that are now exhibiting a higher flat expectation due to the VS being lesioned. Thus, the model proposes that it is the VS input to VTA GABA that gives its expectation signal the temporal specificity that the authors mention in their paper.

## Methods and Materials

### Evaluation of the model

The paradigm used to evaluate the model is a simple CS-US associative learning task and also considers how the expectation cancels out the dopamine peak at the time of the reward. The trial duration is 500 time steps with each time step corresponding to 1ms. The stimulus is presented at the 10^*th*^ time step and is kept switched on till the arrival of the reward at the 400^*th*^ time step (400ms). The reward and the stimulus have by default a magnitude of 1. The number of trials for the entire conditioning to happen was set at 14 trials (i.e. trials required for the learning algorithm to converge).

### Model Description

#### Computational principles

The system-level model is composed of mean-field description of neuronal populations representing distinct, interconnected brain structures. Population dynamics is described by its average firing frequency across time ***U***(*t*), which is taken as the positive part of a membrane potential ***V***(*t*), represented by the following equations:

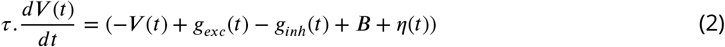

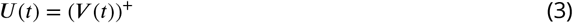

Here *τ* is the time constant of the cell, ***B*** is the baseline firing rate and *η*(*t*) is the additive noise term chosen randomly at each time step from a uniform distribution between −0.01 and 0.01. The incoming afferent synaptic currents *g_exc_* and *g_inh_* represent the weighted sum of excitatory and inhibitory firing rates, respectively, the weight representing the synaptic weights between the populations.

Some of the neuronal populations extract a short-term phasic activity from their incoming inputs, by removing out the tonic component of the input. This is done by the following equations:

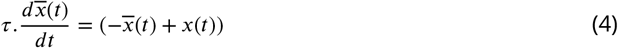

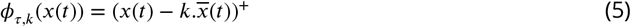

Here 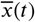 integrates the incoming input *x*(*t*) with a time constant *τ* and thus represents the tonic component of the input, while *ϕ_τ,k_*(*x*(*t*)) represents the positive part of the difference between *x*(*t*) and 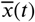. Hence, The constant *k* controls how much of the original input is kept, a *k* value of 0 indicates the entire synaptic input is to be preserved and a *k* value of 1 outputs the phasic component only, i.e. the entire tonic component has been entirely removed.

A Bound function is used when the firing of a population is described with an upper and a lower limit in certain populations.

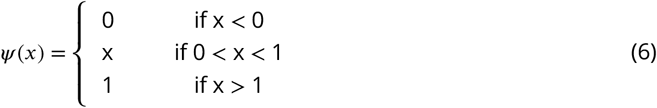

A threshold function is also used in some populations and it outputs 1 when the input exceeds a threshold Γ, 0 otherwise:

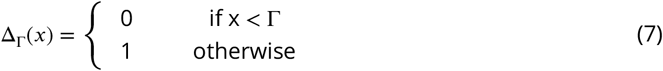

The learning rules defined in the model are based on the Hebbian learning rule and a DA modulated learning rule in the case of BLA like the multiplicative three factor learning rule. The evolution overtime of the weight *w*(*t*) of a synapse between the neuronal population *pre* (presynaptic neurons) and the neuronal population *post* (postsynaptic neurons) is governed by:

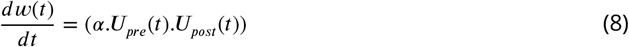

where *w* is the weight term, *α* the learning rate, ***U**_pre_*(*t*) and ***U**_pre_*(*t*) are indicating the firing rates of the presynaptic and postsynaptic neuronal populations, respectively.

### Population definitions

#### Representations of inputs

The sensory inputs of the CS and the reward input of the US are encoded by the inferotemporal cortex (IT) and the lateral hypothalamus (LH), respectively, simply as square wave signals:

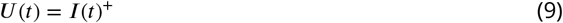

where ***I***(*t*) is an external input resulting either from a stimulus or from a reward.

#### Basolateral Amygdala

The BLA receives inputs about the CS from the IT, the US from the LH, as well as VTA DA output. This allows the BLA to learn to associate the CS with the US, thus providing a magnitude expectation. The equation below is the same equation as in Equation 2 without the inhibitory component and with the presence of a tonic to phasic conversion.

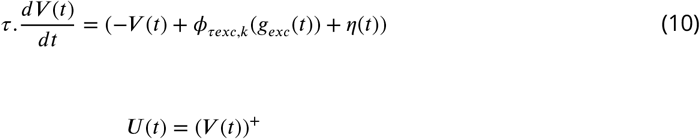

with *τ* = 10ms, *τexc* = 10ms, *k*= 1.

The CS is learned by updating the synaptic weights between IT and BLA and the learning rule is given by:

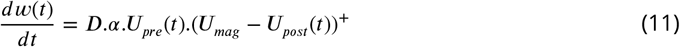

Here ***D*** indicates the presence of the US corresponding to the dopaminergic neuronal modulation from the VTA, *α* is the learning rate equal to 0.003, ***U**_mag_* is the magnitude of LH firing, ***U**_pre_* and ***U**_post_* are the firing rates of presynaptic and postsynpatic neurons, respectively.

#### Central Amygdala

The CE is the output nuclei of the amygdala and it projects to both the PPN nuclei, relaying information from the BLA. The CE projects to the PPN RD neurons that convey US and CS firing to the VTA dopamine neurons and PPN FT neurons that convey reward expectation.

The equations for the membrane potential and the firing rate are the same as Equation 10 and Equation 3, respectively, with *τ* = 20ms, *τexc* = 5ms, *k* = 1.

#### Peduncolopontine nucleus

The PPN has two distinct populations in this model for reward and expectation. The PPN is a heterogeneous structure both in terms of neuronal populations and of responses during classical conditioning (***Okada and Kobayashi, 2009***). Hence, we modeled two distinct subpopulations reflecting the two major classes of responses found experimentally: FT (Fixation Target) population, which activates briefly upon CS or US presentation, and RD (Reward Delivery) population, which display sustained activity during the CS-US interval.

#### PPN RD

The PPN Reward Delivery neurons signal the occurrence of the CS and the US from the CE and the LH, respectively. It also contains a sub-population of inhibitory neurons that inhibits the PPN FT neurons.

The equations for the membrane potential and the firing rate are the same as Equation 10 and Equation 3 respectively, with *τ* = 5ms, *τexc* = 5ms, *k* = 1.

#### PPN FT

The PPN FT neurons encode the magnitude expectation delivered to the VTA GABA neurons. The PPN FT neurons receive information from the CE and are inhibited by the PPN RD neurons. They serve to maintain a constant magnitude that is conveyed to the VTA GABA neurons for final reward prediction error computation. The equations for the membrane potential and the firing rate are the same as Equation 2 without baseline firing and Equation 3 respectively with *τ* = 5ms.

#### Ventral Striatum and OFC

It has long been thought that the Ventral Striatum (VS) is responsible for the reward prediction term in RPE calculation. In our model, the VS encodes the duration of the CS-US interval only. The VS is composed of inhibitory cells, and signals the timing of expected reward to VTA GABA cells through a decrease in activity. The OFC (Orbitofrontal Cortex) relays the presence of the US from the IT to the VS. Then, a simplified timing model comprising a negative integrator similar to the timing algorithm in ***Rivest and Bengio*** (***2011***) signals the interval duration through a slowly decreasing activity until the expected timing of the US. To do so, the integrator here has an amplitude of 1 at the beginning of the trial and after weight updating, decreases its firing to 0 at the precise time of reward delivery. In this framework, learning the CS-US interval consists in adjusting the slope of the slowly-decreasing activity.

#### Mechanism of timing

The timing mechanism in the VS transforms a phasic excitatory input into a decreasing sustained activity, which slope depends upon the weights.

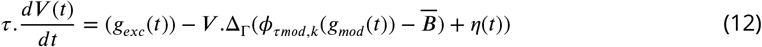

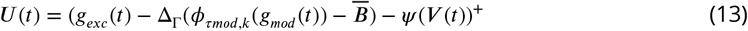

with *τ* = 1ms, *τmod* = 5ms, *k* = 1, Γ = 6 and 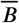 is the baseline firing rate from VTA dopamine to VS. Γ ensures a minimum threshold to be achieved for the VTA dopamine phasic firing to enable modulation. *ψ*() is a bounded function.

As described in figure 6, weight is updated after each iteration according to the following rule:

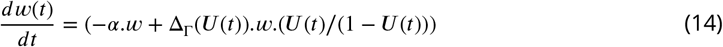

where *α* is the learning rate equal to 0.4 The first term decreases the weights based on *α* and the weights keep decreasing until the bound is reached when Δ_Γ_(***U***(*t*)) becomes greater than 0 at the time of the reward. The correcting update is the second term of the weight updating and the slope is increased with a weight increase encoding the duration of the interval.

**Figure 6.**
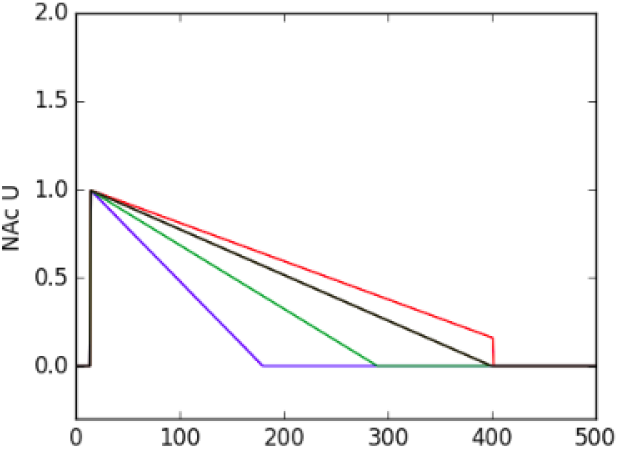
The slope is decreased at every iteration until it exceeds the duration (the red line) enabling exact correction of the weight encoding the duration to be found (the black line). The colors indicate the progressive iterations

It should be noted that the model postulates that the learning of time happens before the learning of value of the stimulus, i.e., its magnitude.

#### VTA

The VTA comprises two major neuronal populations, dopaminergic (DA) and gabaergic (GABA), glutamatergic cells representing less than 3 percent of VTA cells. VTA GABA neurons locally inhibit VTA DA neurons and participate in the computation of reward prediction error in VTA DA cells (***Cohen et al., 2012***). More precisely, VTA GABA neurons display a sustained, slowly-increasing ramping activity during the CS-US interval (***Eshel et al., 2015***) but only significantly affect phasic DA activity (i.e. the RPE) rather tonic DA activity during the interval. We thus modeled the two populations from the VTA as follows.

#### VTA Dopamine

The VTA dopaminergic (DA) neurons receive excitatory inputs from the PPN RD population, which conveys actual reward and reward prediction from the amygdala; and inhibitory inputs from the VTA GABA cells, which signal reward expectation. The difference between these excitatory and inhibitory inputs constitutes the reward prediction error (RPE) (***Sutton and Barto, 1998***) (***Glimcher, 2011***). VTA DA neurons broadcast this RPE to the system. During learning, VTA DA neurons initially fire upon US reward delivery. This US activity progressively gets canceled by VTA GABA signaling the reward expectation, and at the same time, phasic firing upon CS arrival develops with learning.

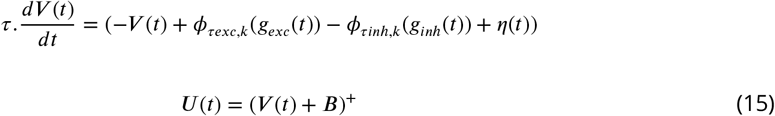

With *τ* = 5ms, *τexc* = 5ms, *k* = 1 and ***B*** is the baseline firing rate of the VTA Dopamine equal to 0.2

#### VTA GABA

VTA GABA neurons combine inputs from the VS, which encodes the expected time of reward, and from the PPN, which signals expected reward magnitude. VTA GABA neurons thus encode reward expectation and inhibit VTA DA neurons. The equation for membrane potential is the same as in Equation 2 without baseline firing and population dynamics follows Equation 3 with *τ* = 20ms.

This model is implemented in Python, and uses the DANA library for neuronal computation (***Rougier and Fix, 2012***). Description of all the other model parameters is detailed in Table 1. The model can be accessed in the following link: https://github.com/palladiun/Pavlovian-Conditioning-VTA-GABA

**Table 1.** Table describing network architecture and parameters used in activation and learning rules.

| Architectural parameters |  |  |
| --- | --- | --- |
| Parameter | Meaning | Value |
| US input_size | size of input vectors from LH | 1 |
| CS input_size | size of input vectors from IT | 4 |
| VTA Dopamine_size | number of neurons in VTA Dopamine | 10 |
| VTA GABA_size | number of neurons in VTA GABA | 5 |
| BLA_size | number of neurons in BLA | 1 |
| CE_size | number of neurons in CE | 1 |
| OFC_size | number of neurons in OFC | 1 |
| PPN RD_size | number of neurons in PPN RD | 4 |
| PPN FT_size | number of neurons in PPN FT | 4 |
| PPN Magnitude_size | number of neurons in PPN Magnitude | 4 |
| Equation parameters |  |  |
| BLA_CE | constant weights from BLA to CE | 0.15 |
| LH_PPN_RD | constant weights from LH to PPN_RD | 1.2 |
| LH_BLA | constant weights from LH to BLA | 1 |
| IT_OFC | constant weights from IT to OFC | .25 |
| CE_PPN_RD | constant weights from CE to PPN_RD | 2 |
| CE_PPN_Mag | constant weights from CE to PPN_Mag | .3 |
| PPN_RD_PPN_Mag | constant weights from PPN_RD to PPN_Mag | 0.8 |
| PPN_RD_VTA_Dop | constant weights from PPN_RD to VTA_Dop | 1 |
| PPN_Mag_PPN_Rel | constant weights from PPN_Mag to PPN_Rel | 0.2 |
| VS_PPN_Rel | constant weights from VS to PPN_Rel | 1 |
| PPN_Rel_VTA_GABA | constant weights from PPN_Rel to VTA_GABA | 0.25 |
| VTA_Dopamine_BLA | constant weights from VTA_Dopamine to BLA | 1 |
| VTA_Dopamine_VS | constant weights from VTA_Dopamine to VS | 1 |
| VTA_GABA_VTA_Dopamine | constant weights between VTA_GABA and VTA_Dopamine | 0.2 |
| OFC_VS | initial weights between OFC and VS | 0.006 |
| IT_BLA | initial weights between OFC and VS | 0.01 |

## Acknowledgments

We would like to acknowledge the following grants which have been a major support to this research.

- Indo-French CEFIPRA Grant for the project Basal Ganglia at Large (No. DST-INRIA2013-02/Basal Ganglia dated 13-09-2014)
- Internships programme at INRIA, 6 month Internship with Team Mnemosyne at INRIA Bordeaux - Sud-Ouest

We also thank Maxime Carrere for helpful discussions.

## References

Bocklisch C, Pascoli V, Wong JC, House DR, Yvon C, De Roo M, Tan KR, Lüscher C. Cocaine disinhibits dopamine neurons by potentiation of GABA transmission in the ventral tegmental area. Science. 2013; 341(6153):1521–1525.

Carmichael S, Price JL. Sensory and premotor connections of the orbital and medial prefrontal cortex of macaque monkeys. Journal of Comparative Neurology. 1995; 363(4):642–664.

Chuhma N, Tanaka KF, Hen R, Rayport S. Functional connectome of the striatal medium spiny neuron. Journal of Neuroscience. 2011; 31(4):1183–1192.

Cohen JY, Haesler S, Vong L, Lowell BB, Uchida N. Neuron-type specific signals for reward and punishment in the ventral tegmental area. nature. 2012; 482(7383):85.

Dayan P. Improving generalization for temporal difference learning: The successor representation. Neural Computation. 1993; 5(4):613–624.

Eshel N, Bukwich M, Rao V, Hemmelder V, Tian J, Uchida N. Arithmetic and local circuitry underlying dopamine prediction errors. Nature. 2015; 525(7568):243.

Eshel N, Tian J, Bukwich M, Uchida N. Dopamine neurons share common response function for reward prediction error. Nature neuroscience. 2016; 19(3):479.

Fiorillo CD, Newsome WT, Schultz W. The temporal precision of reward prediction in dopamine neurons. Nature neuroscience. 2008; 11(8):966–973.

Fiorillo CD, Tobler PN, Schultz W. Discrete coding of reward probability and uncertainty by dopamine neurons. Science. 2003; 299(5614):1898–1902.

Fiorillo CD, Tobler PN, Schultz W. Evidence that the delay-period activity of dopamine neurons corresponds to reward uncertainty rather than backpropagating TD errors. Behavioral and brain Functions. 2005; 1(1):7.

Glimcher PW. Understanding dopamine and reinforcement learning: the dopamine reward prediction error hypothesis. Proceedings of the National Academy of Sciences. 2011; 108(Supplement 3):15647–15654.

Hollerman JR, Schultz W. Dopamine neurons report an error in the temporal prediction of reward during learning. Nature neuroscience. 1998; 1(4):304.

Hong S, Hikosaka O. Pedunculopontine tegmental nucleus neurons provide reward, sensorimotor, and alerting signals to midbrain dopamine neurons. Neuroscience. 2014; 282:139–155.

Janak PH, Tye KM. From circuits to behaviour in the amygdala. Nature. 2015; 517(7534):284.

Joel D, Niv Y, Ruppin E. Actor-critic models of the basal ganglia: New anatomical and computational perspectives. Neural networks. 2002; 15(4-6):535–547.

Keiflin R, Janak PH. Dopamine prediction errors in reward learning and addiction: from theory to neural circuitry. Neuron. 2015; 88(2):247–263.

Klein-Flügge MC, Hunt LT, Bach DR, Dolan RJ, Behrens TE. Dissociable reward and timing signals in human midbrain and ventral striatum. Neuron. 2011; 72(4):654–664.

Kobayashi Y, Okada KI. Reward prediction error computation in the pedunculopontine tegmental nucleus neurons. Annals of the New York Academy of Sciences. 2007; 1104(1):310–323.

Lokwan S, Overton P, Berry M, Clark D. Stimulation of the pedunculopontine tegmental nucleus in the rat produces burst firing in A9 dopaminergic neurons. Neuroscience. 1999; 92(1):245–254.

Niv Y, Duff MO, Dayan P. Dopamine, uncertainty and TD learning. Behavioral and brain Functions. 2005; 1(1):6.

Okada KI, Kobayashi Y. Characterization of oculomotor and visual activities in the primate pedunculopontine tegmental nucleus during visually guided saccade tasks. European Journal of Neuroscience. 2009; 30(11):2211–2223.

Okada Ki, Kobayashi Y. Reward prediction-related increases and decreases in tonic neuronal activity of the pedunculopontine tegmental nucleus. Frontiers in integrative neuroscience. 2013; 7.

O’reilly RC, Frank MJ, Hazy TE, Watz B. PVLV: the primary value and learned value Pavlovian learning algorithm. Behavioral neuroscience. 2007; 121(1):31.

Pan WX, Schmidt R, Wickens JR, Hyland BI. Dopamine cells respond to predicted events during classical conditioning: evidence for eligibility traces in the reward-learning network. Journal of Neuroscience. 2005; 25(26):6235–6242.

Pavlov IP. Conditional reflexes: An investigation of the physiological activity of the cerebral cortex. H. Milford; 1927.

Rivest F, Bengio Y. Adaptive drift-diffusion process to learn time intervals. arXiv preprint arXiv:11032382. 2011;.

Rougier NP, Fix J. DANA: distributed numerical and adaptive modelling framework. Network: Computation in Neural Systems. 2012; 23(4):237–253.

Sadacca BF, Jones JL, Schoenbaum G. Midbrain dopamine neurons compute inferred and cached value prediction errors in a common framework. Elife. 2016; 5:e13665.

Sah P, Faber EL, De Armentia ML, Power J. The amygdaloid complex: anatomy and physiology. Physiological reviews. 2003; 83(3):803–834.

Schultz W, Dayan P, Montague PR. A neural substrate of prediction and reward. Science. 1997; 275(5306):1593–1599.

Semba K, Fibiger HC. Afferent connections of the laterodorsal and the pedunculopontine tegmental nuclei in the rat: a retro-and antero-grade transport and immunohistochemical study. Journal of Comparative Neurology. 1992; 323(3):387–410.

Sutton RS, Barto AG. Reinforcement learning: An introduction, vol. 1. MIT press Cambridge; 1998.

Takahashi YK, Langdon AJ, Niv Y, Schoenbaum G. Temporal specificity of reward prediction errors signaled by putative dopamine neurons in rat VTA depends on ventral striatum. Neuron. 2016; 91(1):182–193.

Usuda I, Tanaka K, Chiba T. Efferent projections of the nucleus accumbens in the rat with special reference to subdivision of the nucleus: biotinylated dextran amine study. Brain research. 1998; 797(1):73–93.

Vitay J, Hamker FH. Timing and expectation of reward: a neuro-computational model of the afferents to the ventral tegmental area. Frontiers in neurorobotics. 2014; 8.

Xia Y, Driscoll JR, Wilbrecht L, Margolis EB, Fields HL, Hjelmstad GO. Nucleus accumbens medium spiny neurons target non-dopaminergic neurons in the ventral tegmental area. Journal of Neuroscience. 2011; 31(21):7811–7816.

Yau HJ, Wang DV, Tsou JH, Chuang YF, Chen BT, Deisseroth K, Ikemoto S, Bonci A. Pontomesencephalic tegmental afferents to VTA non-dopamine neurons are necessary for appetitive Pavlovian learning. Cell reports. 2016; 16(10):2699–2710.

Yoo JH, Zell V, Wu J, Punta C, Ramajayam N, Shen X, Faget L, Lilascharoen V, Lim BK, Hnasko TS. Activation of pedunculopontine glutamate neurons is reinforcing. Journal of Neuroscience. 2017; 37(1):38–46.

